# Evidence for endemism and local adaptation in Antarctic soil bacteria

**DOI:** 10.64898/2026.05.01.722257

**Authors:** Nicholas B. Dragone, Mary K. Childress, Noah Mendez, Jordan Galletta, Caihong Vanderburgh, Clifton P. Bueno de Mesquita, Kristen M. DeAngelis, C. Alisha Quandt, Pok Man Leung, Chris Greening, Byron J. Adams, Noah Fierer

**Author notes:** Corresponding authors: Nicholas B. Dragone and Noah Fierer. **Author Contributions:** Conceptualization: NBD, NF, BJA, CAQ Methodology: NBD, NF, BJA, CAQ, MKC, CPB, CG, PML Investigation: NBD, MKC, NM, JG, CV, CPB, Visualization: NBD, MKC, NF, NM Funding acquisition: NBD, NF, BJA, CAQ, CG, PML Project administration: NBD, BJA, CAQ Supervision: NF, BJA, CAQ, CG, KMD Writing – original draft: NBD, NF Writing – review & editing: NBD, MKC, NM, JG, CV, CPB, KMD, CAQ, PML, CG, BJA, NF. **Competing Interest Statement:** Authors declare that they have no competing interests.

## Abstract

Antarctic soils represent one of the more extreme environments for microbial life on Earth, yet they harbor heterogeneous and diverse microbial communities. Biologists have long hypothesized that Antarctic microorganisms are unique from those found on other continents due to the extreme geographic isolation and the cold, dry, and challenging conditions typical of Antarctica. To test this hypothesis, we focused on a cosmopolitan bacterial genus, *Arthrobacter*, that is widely distributed across global soils. We first profiled a global metagenomic dataset from both Antarctic and non-Antarctic surface soils to quantify the distributions of *Arthrobacter* strains. Despite high strain-level diversity, 90% of the strains found in the Antarctic soils were only found on the continent. We then used cultivation-based phenotypic analyses and strain-level genomic comparisons to assess how Antarctic strains and non-Antarctic strains differ in their traits and environmental preferences. Not only did we find evidence of endemism, but Antarctic *Arthrobacter* also have genomic characteristics and environmental tolerances that suggest they are uniquely adapted to Antarctic conditions.

**Significance Statement:** Antarctic soils are among the most extreme environments on Earth, yet they host diverse microbial communities whose adaptations are poorly understood. To test whether Antarctic microbes are distinct from those elsewhere, we examined *Arthrobacter*, a bacterial genus common in soils worldwide. Analysis of global metagenomic data revealed *Arthrobacter* strains in Antarctic soils are found exclusively on the continent. Cultivation experiments and comparative genomics further showed that Antarctic strains differ from non⍰Antarctic relatives in genomic features and environmental tolerances. Together, these results demonstrate that geographically isolated and extreme conditions can drive microbial endemism and local adaptation, even within globally distributed bacterial lineages.

## Main Text

### Introduction

Endemic species are species that are exclusively found in a specific geographic location or region (1, 2). While the most famous examples include charismatic megafauna on isolated islands – Darwin’s finches in the Galapagos (3), koalas in Australia (4), kiwi in Aotearoa New Zealand (5), endemism is not restricted to specific geographic scales and can occur across many taxonomic groups. For example, numerous carnivorous plants are endemic to the Guiana highlands of South America (6) and large freshwater lakes in East Africa have long been known to harbor endemic fish species (7). Indeed, endemism can arise from a number of factors, most notably geographic isolation or the uniqueness of environmental conditions (including historical conditions) (1, 8, 9).

Endemic plants, animals, insects, and even macro-fungi have been described, but are there endemic microorganisms? Given their potential capacity for long-distance dispersal (10–12), we might expect that most microorganisms are cosmopolitan (9). However, evidence of endemism has been described in certain microbial groups, most notably in host-associated organisms and extreme environments. Certain *Metschnikowia* yeasts have been described as endemic to Hawaiian beetles (13), endemic diatoms have been identified in Antarctic lakes (14) and cultivation-independent approaches were used to show that the bacterial genus *Venenivibrio* (phylum Aquificota) is restricted to geothermal springs within Aotearoa New Zealand (15). These examples typify most reports of microbial endemism where evolutionary divergence is driven by geographic isolation or an extremely restricted ecological niche (e.g., a host only found in one place or specific geothermal conditions). It seems unlikely that we would find endemic microorganisms in a more globally ubiquitous environment like soil that harbors highly diverse microbial communities and high micro-site heterogeneity in environmental conditions (16–18).

If endemic soil microorganisms do exist, we might expect to find them in Antarctica (19). Soils on the Antarctic continent are geographically isolated from soil on other continents and experience conditions distinct from those found nearly anywhere else on Earth (20, 21), factors that can lead to endemism (1, 2, 9). For one, significant barriers limit organismal dispersal into and out of Antarctica, including the atmospheric circumpolar vortex, which limits aeolian dispersal into and out of the Antarctic continent (22–25). Antarctic soils are also some of the coldest and driest soils on Earth (26), spanning gradients in climate and edaphic characteristics (27–29) which can impact the structure and functional attributes of Antarctic soil microbial communities (29–32). Given that many of the soils found on the Antarctic continent have likely remained ice-free for thousands of years (33), with their microbial communities potentially developing in isolation from the rest of the world, certain soil microorganisms may be endemic to the continent, but this hypothesis remains largely untested.

What characteristics and genomic attributes would endemic Antarctic soil microbes have? Given the harsh environmental conditions typical of most Antarctic soils, we would expect that Antarctic soil microorganisms are locally adapted to the extreme conditions experienced in Antarctica, including possible adaptations to survive extremely cold, arid, and nutrient-poor conditions characteristic of soils from the southern continent (20, 21). Such adaptations could include genes associated with cold tolerance, desiccation tolerance, oligotrophy (34), trace gas metabolism (35) or other metabolic strategies that have been hypothesized to be associated with Antarctic bacteria (29, 36).

Here, we searched for signs of endemism and local adaptation in Antarctic soil microorganisms by using cultivation-independent metagenomic sequencing. To do this, we assessed strain-level biogeographic distributions of the bacterial genus *Arthrobacter* in 126 soils from across the Antarctic continent and 472 non-Antarctic soils from other regions across the globe (including soils from other cold and dry systems). We focus on this genus of Gram-positive, obligate aerobes as *Arthrobacter* are relatively abundant in soils across the globe (37, 38), especially in cold and dry environments. Moreover, members of this genus are relatively ubiquitous in Antarctic soils (30, 36, 37) and are readily culturable. We first tested the hypothesis that Antarctic *Arthrobacter* strains are distinct from those found in non-Antarctic soils from across the globe. We then determined whether Antarctic *Arthrobacter* strains also have distinct genomic attributes and environmental tolerances, reflecting the unique adaptations of members of this genus to survive harsh Antarctic conditions. To do so, we paired the biogeographic analyses with genomic analyses of 171 soil *Arthrobacter* strains and detailed phenotypic analyses of 53 *Arthrobacter* isolates from both Antarctic and non-Antarctic soils to quantify in vitro growth rates across gradients in temperature and water availability.

## Results and Discussion

### Evidence for endemicity

Endemic taxa are, by definition, restricted in their geographic distribution and found only in certain regions (1, 8, 9). In our case, for *Arthrobacter* in Antarctic soils to be endemic, particular strains or species should only be found in Antarctica and not in other soils from across the globe. To test this hypothesis, we profiled 598 metagenomes in total, 126 from Antarctic soils and 472 from non-Antarctic soils that spanned a broad range in climates and ecosystem types, Figure 1). We then searched for the presence of any *Arthrobacter* strain from our genome reference database that included 568 unique *Arthrobacter* strains (see Methods). While we recognize that there are likely *Arthrobacter* strains not represented in our analysis, our reference database spans the full breadth of known diversity across the genus and includes both cultivated taxa as well as metagenome-assembled genomes.

**Figure 1.**
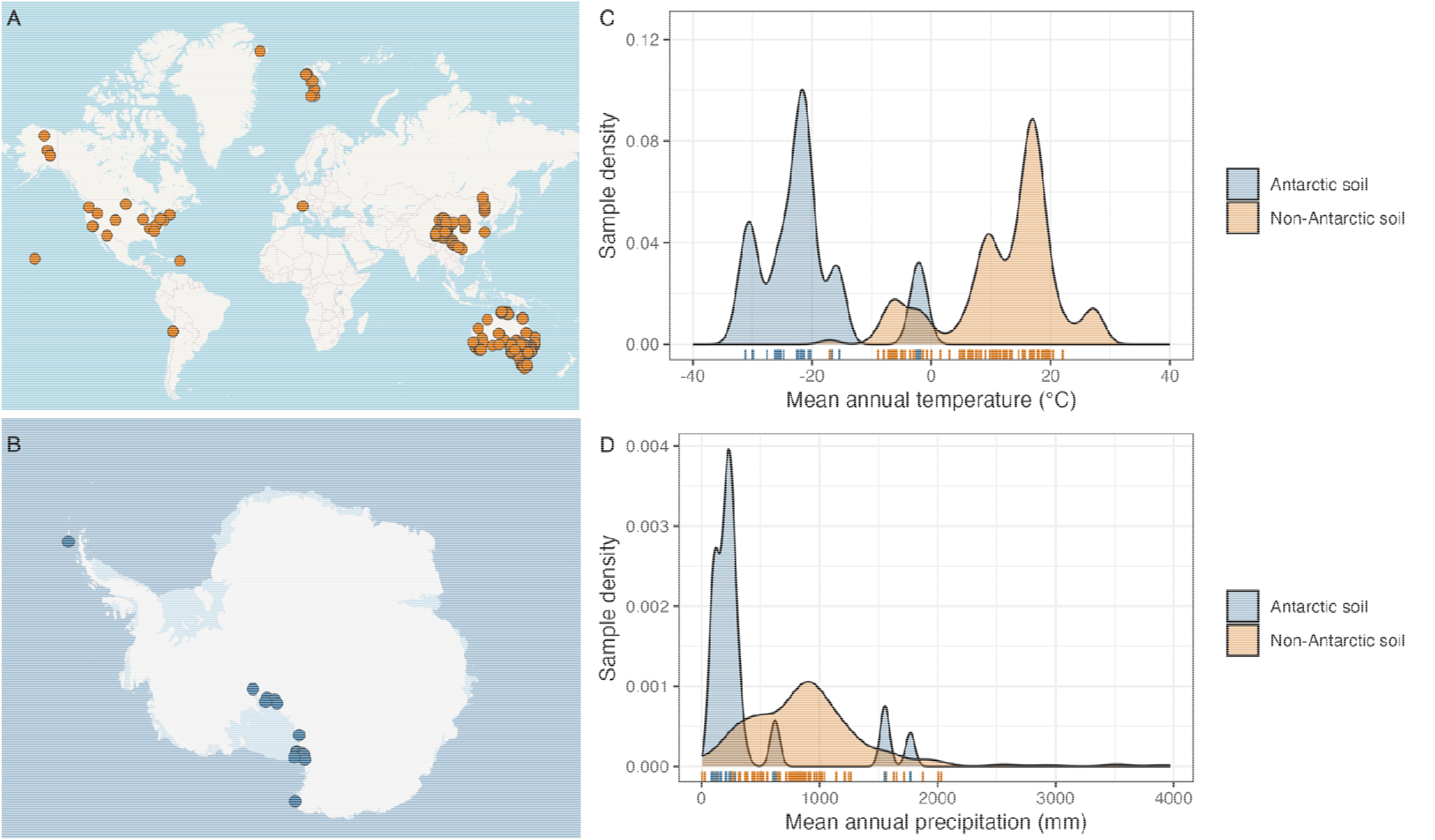
**A)** Map displaying the collection locations of the 472 non-Antarctic soil metagenomes used for metagenomic profiling. **B)** Map displaying the locations of the 17 ice-free regions from which the 126 Antarctic soil metagenomes were collected. **C)** The distribution of mean annual temperatures (MAT) across the Antarctic soil metagenomes (n = 126) and non-Antarctic soil metagenomes (n = 472) included in this study. **D)** The distribution of mean annual precipitation values (MAP) across the Antarctic and non-Antarctic soil metagenomes included in this study. Y-axis values show calculated frequency density of each group. The skirt plots indicate the MAT and MAP of the 50 Antarctic soils and 105 non-Antarctic soils in which one or more *Arthrobacter* strains were detected.

Across the 126 Antarctic soil metagenomes profiled, at least one *Arthrobacter* strain (mean of 2 per sample, range 1 – 7 strains) was detected in 50 of the metagenomes (40%, Figure 2). Of the 472 non-Antarctic soil metagenomes profiled, an *Arthrobacter* strain was detected in 105 metagenomes (22.2%, mean of 2.3 per sample, range 1 – 12). More generally, we found *Arthrobacter* to be reasonably ubiquitous in Antarctic soils, as has been reported previously and were identified across the range of climatic conditions (Figure 1). In total, we detected 36 *Arthrobacter* strains in Antarctic soils and 111 strains in non-Antarctic soils. Of these strains, only four strains were detected in both Antarctic and non-Antarctic soils, significantly lower than expected under the null model (p < 0.001) (Figure S1) which suggests that this is unlikely to be a random occurrence (Hypergeometric probability, P < 0.0001). These results are supported by the significant differences identified between the Antarctic and non-Antarctic *Arthrobacter* communities, as determined by permutational ANOVAs (unweighted Unifrac distances, F=13.527, p=0.001; Jaccard distances, F = 3.46, p = 0.001). The four overlapping strains, which included psychrophilic strains (e.g. *A. psychrotolerans*, Figure 2), were only found in non-Antarctic soils from relatively cold and dry systems including the Arctic (Svalbard), the Swiss Alps, and the Tibetan Plateau.

**Figure 2.**
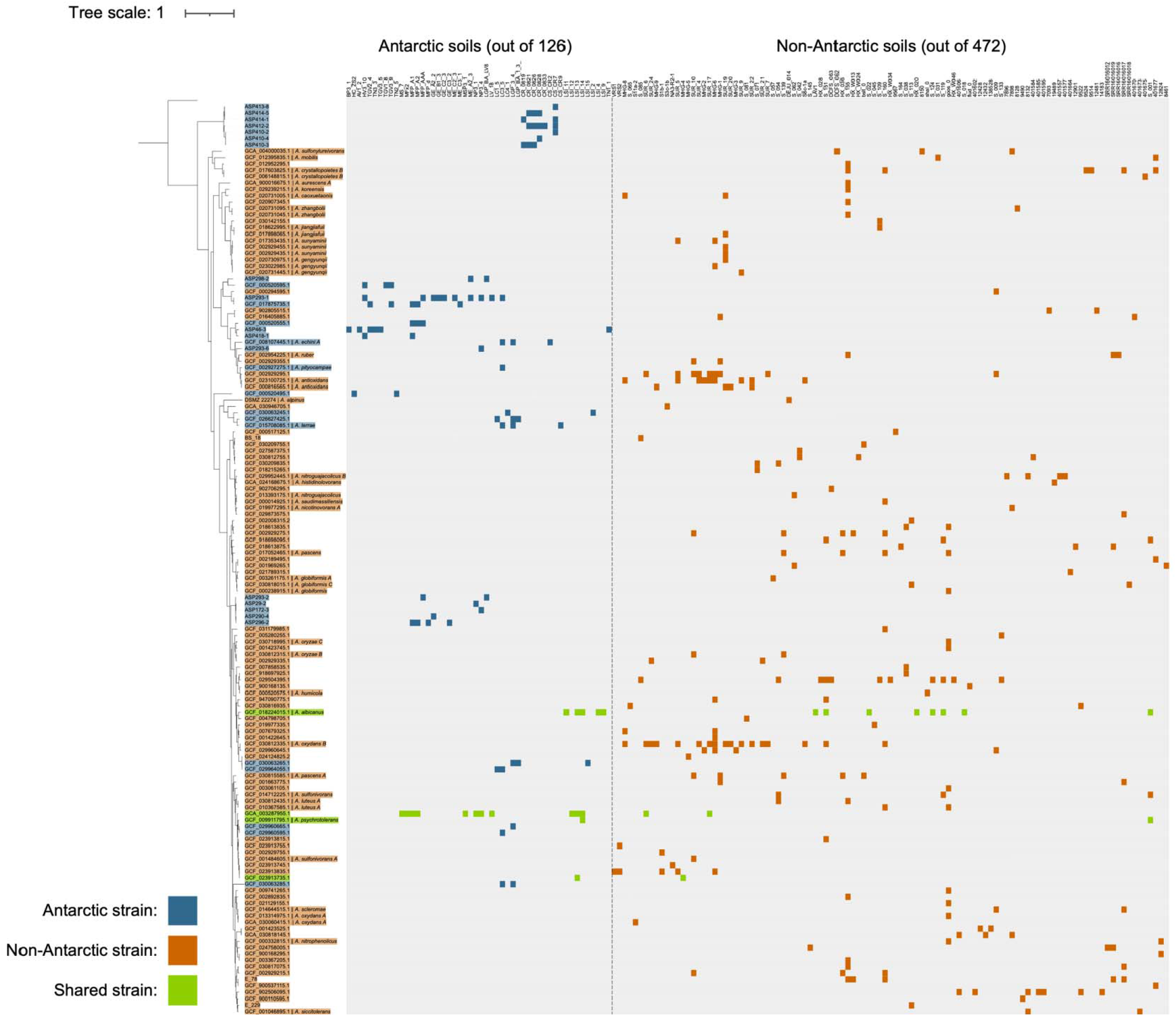
*Arthrobacter* strains identified in Antarctic and non-Antarctic soil metagenomes. Phylogenetic tree represents the 32 Antarctic, 107 non-Antarctic, and 4 shared *Arthrobacter* strains identified via metagenomic profiling. Strains are labeled either with the genome ID or culture ID – with species identity provided if available. Strain labels are highlighted in blue if they were detected in only Antarctic soils, orange if only detected in non-Antarctic soils, and green if shared across the two sample categories. The heatmap to the right of the tree displays the 50 Antarctic soil metagenomes and 105 non-Antarctic soil metagenomes that were identified as containing at least one Arthrobacter strain. If a strain was identified in a metagenome it is highlighted with the colors described previously. Metagenomes for both sample categories are ordered from lowest mean annual temperature (MAT) to highest (left to right).

While definitive proof of endemism is difficult to obtain as we can never rule out the possibility that any taxon might be found in some soil somewhere, our findings suggest that 90% of the *Arthrobacter* strains detected in Antarctic soils (32/36) were not found elsewhere and are thus likely endemic to the Antarctic continent (Figure 2). Given our sequencing depth and the relative abundance thresholds required for strain detection, our estimates of endemicity are conservative with respect to low-abundance cosmopolitan strains. However, Antarctic strains do not fall into a single distinct phylogenetic group. Instead, the 32 strains that were exclusively detected in Antarctic soils and the 107 strains that were only detected in the non-Antarctic soils are distributed across the *Arthrobacter* phylogenetic tree (Figure 2). Nevertheless, the phylogenetic diversity (PD) of strains found in Antarctica (Faith’s PD = 4.89) is lower than that of the non-Antarctic strains (Faith’s PD = 8.02) which suggests that Antarctic *Arthrobacter* are less phylogenetically diverse, possibly reflecting a more limited, and shared, Antarctic evolutionary history.

The phylogenetic structure we observe is consistent with deep-time history of continental isolation and evolutionary filtering in Antarctica. Following the establishment of circumpolar circulation during the Oligocene–Miocene, together with the onset of major Antarctic glaciation at the Eocene–Oligocene transition (∼34 Ma), Antarctica has remained strongly thermally and climatically isolated for tens of millions of years (39, 40). This prolonged isolation, coupled with repeated glacial–interglacial cycles and the persistence of ice-free refugia and exceptionally stable landscapes in the in Transantarctic Mountains since the mid-Miocene (∼14-13Ma) (41, 42), would have constrained dispersal while promoting in situ diversification and regional persistence of microbial lineages. Under such a regime, repeated colonization events followed by long-term ecological filtering and demographic bottlenecks could yield the pattern observed here: phylogenetically dispersed Antarctic strains with reduced overall diversity relative to the global pool. In this context, the strain-level endemism detected in *Arthrobacter* likely reflects not only contemporary ecological selection, but also the cumulative evolutionary response to millions of years of isolation.

### Evidence of local adaptation

We expected that endemic Antarctic organisms would be locally adapted to Antarctic conditions and have different environmental tolerances from closely related organisms found in other environments due to the unique conditions found on the continent. Thus, we next sought to compare the optimal growth temperatures and desiccation tolerances of our Antarctic and non-Antarctic *Arthrobacter* strains, testing the hypothesis that Antarctic *Arthrobacter* are adapted to the cold temperatures and hyper-arid conditions typical of inland soils on the continent. While we recognize that there are many environmental stressors faced by Antarctic soil microorganisms (20, 43, 44), we focused on growth responses across gradients in temperature and water availability because these factors are most often associated with variation in the overall structure of Antarctic soil communities (29, 30, 32, 45, 46).

To compare the temperature optima of Antarctic and non-Antarctic *Arthrobacter*, we grew 28 Antarctic *Arthrobacter* isolates and 25 non-Antarctic *Arthrobacter* isolates at five different temperatures (4°, 11°, 18°, 25°, 32°C) under conditions of maximum water availability. Representative isolates from each group grew across all temperatures tested. However, the Antarctic *Arthrobacter* isolates had a significantly lower temperature optima than non-Antarctic isolates (Mann-Whitney U, p < 0.001, Figure 3). The mean temperature optimum of Antarctic isolates was 16.7°C (range 12 -24°C) while the mean temperature optimum of non-Antarctic isolates was 27.6°C (range 18 -32°). Antarctic isolates also grew, on average, >300% slower than non-Antarctic isolates at the optimal growth temperature for each strain (Figure 3).

**Figure 3.**
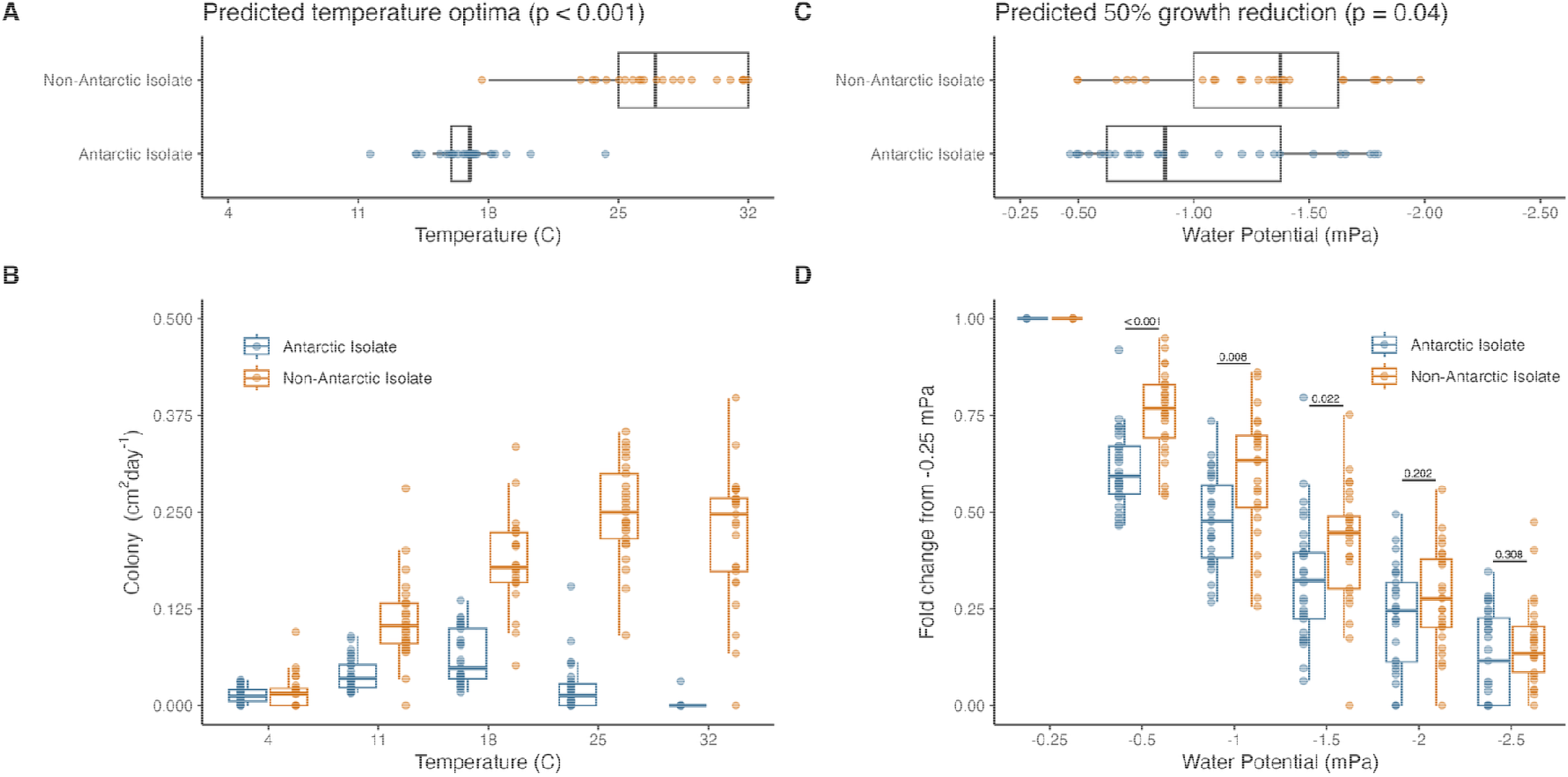
Environmental tolerances of Antarctic (n=28) and non-Antarctic (n=25) *Arthrobacter* isolates. **A)** Predicted temperature optima of *Arthrobacter* isolates based on generalized additive models (GAMs), calculated after seven days of growth on media with maximum water availability. Each point represents modeled Topt for a single isolate. **B)** Colony growth at five different temperatures after seven days. Each dot represents the average growth across the four replicates for each isolate at each of the five time points. **C)** Predicted water potential at which growth of each isolate is reduced to 50% of maximum. Each dot represents the water potential modeled for one isolate. **D)** Fold change from growth at -0.25 MPa water potential (0% PEG) across our desiccation gradient. Each point displayed represents the average across the four replicates for each isolate. Significance of the difference between two groups is displayed above each water potential.

Our finding that Antarctic strains have significantly lower optimal growth temperatures and lower growth rates at optimal growth temperature than non-Antarctic strains is not surprising: to persist and be active in Antarctica, microorganisms must be able to tolerate and grow at extremely low temperatures, tolerances often associated with slower growth rates (44). What is notable is that the optimal growth temperatures of our Antarctic isolates (12 - 24oC) are far higher than the mean annual temperatures experienced by the soils from which our strains were isolated (range: -1.9 to -37.6 °C, see ref (30)). Why would Antarctic *Arthrobacter* not have a growth temperature optimum that are more closely aligned with *in situ* conditions? We suspect this is due to the pronounced seasonal variation in Antarctic soil temperature regimes. During the polar summer months, soil surface temperatures can be well above freezing, especially on exposed soil surfaces that can receive nearly 24h of direct sunlight (47). It is possible that Antarctic *Arthrobacter* strains are “opportunitrophs” (48) that remain largely inactive - or grow extremely slowly - for most of the year, and only become active when conditions are favorable, a growth strategy that has been described (or predicted) in other cold and dry environments (49, 50).

We also measured growth responses of all 53 strains across gradients in water availability (-0.25 to -2.5 mPa) at near-optimal growth temperatures for each strain (see Methods). In contrast to expectations, we did not find any evidence that Antarctic soil *Arthrobacter* are more tolerant to desiccation, i.e., able to grow better at lower water (matric) potentials than isolates from non-Antarctic soils. In fact, our Antarctic *Arthrobacter* strains were less desiccation tolerant than non-Antarctic *Arthrobacter* strains (Mann-Whitney U, p = 0.04, Figure 3C) with growth reduced by 50% at a water potential of -1.0 mPa (-0.5 – -1.75 mPa) for Antarctic strains and -1.26 mPa (-0.5 – -2.0 mPa) for non-Antarctic isolates. However, this difference was only observed at intermediate water potentials (-0.5 – 1.5 MPa water potential). At lower water potentials (below -1.5 MPa), there were no significant differences in average growth rates between Antarctic and non-Antarctic isolates (Figure 3D). However, we note that our in vitro assays did not include water potentials below -2.5 MPa and Antarctic soils frequently experience water potentials lower than -3.5 MPa (20, 51). Moreover, our Antarctic isolates exhibited less reduction of growth at more negative water potential (linear regression: y = -0.34x + 0.9, R2 = 0.74) than non-Antarctic isolates (linear regression: y = -0.35x + 0.99, R^2^ = 0.81). This suggests that, although we did not find evidence that Antarctic *Arthrobacter* are more desiccation tolerant than non-Antarctic *Arthrobacter*, Antarctic strains may be able to grow at more extreme levels of desiccation, a hypothesis that should be tested in the future.

### Comparative Genomics

We next used strain-level genomic analyses to build on the cultivation-dependent analyses described above and more broadly determine if the Antarctic *Arthrobacter* strains have unique adaptations for growth and persistence in Antarctic soils. These genomic analyses were conducted by comparing the genomes of 171 *Arthrobacter* strains – 42 Antarctic strains and 129 non-Antarctic strains (Figure S2), to identify genomic attributes and genome-derived traits that differentiate exclusively Antarctic *Arthrobacter* strains from those detected in non-Antarctic soils.

First, our genome-derived estimates of optimal growth temperature largely supported our culture-dependent measurements: Antarctic *Arthrobacter* isolates had a significantly lower estimated temperature optima than non-Antarctic isolates (Mann-Whitney U, p < 0.001, Figure 4A). The average estimated Topt of Antarctic strains was 19.5°C (range 14.5 - 24.6°C) while the average estimated Topt of non-Antarctic strains was 23.4°C (range 17.6 – 40.3°C). While not identical to the measurements from our cultivation-dependent assays, genome-predicted Topt and measured Topt were significantly correlated for the 53 strains that overlapped across the two datasets (Pearson’s r = 0.55, p < 0.001). Moreover, the genome-derived estimates suggest that Antarctic strains have significantly lower minimum growth temperature, within the ranges of soil temperatures commonly measured in polar summer, than non-Antarctic strains (52) (Mann-Whitney U, p < 0.001, Figure S3B) which supports the “opportunitrophs” hypothesis.

**Figure 4.**
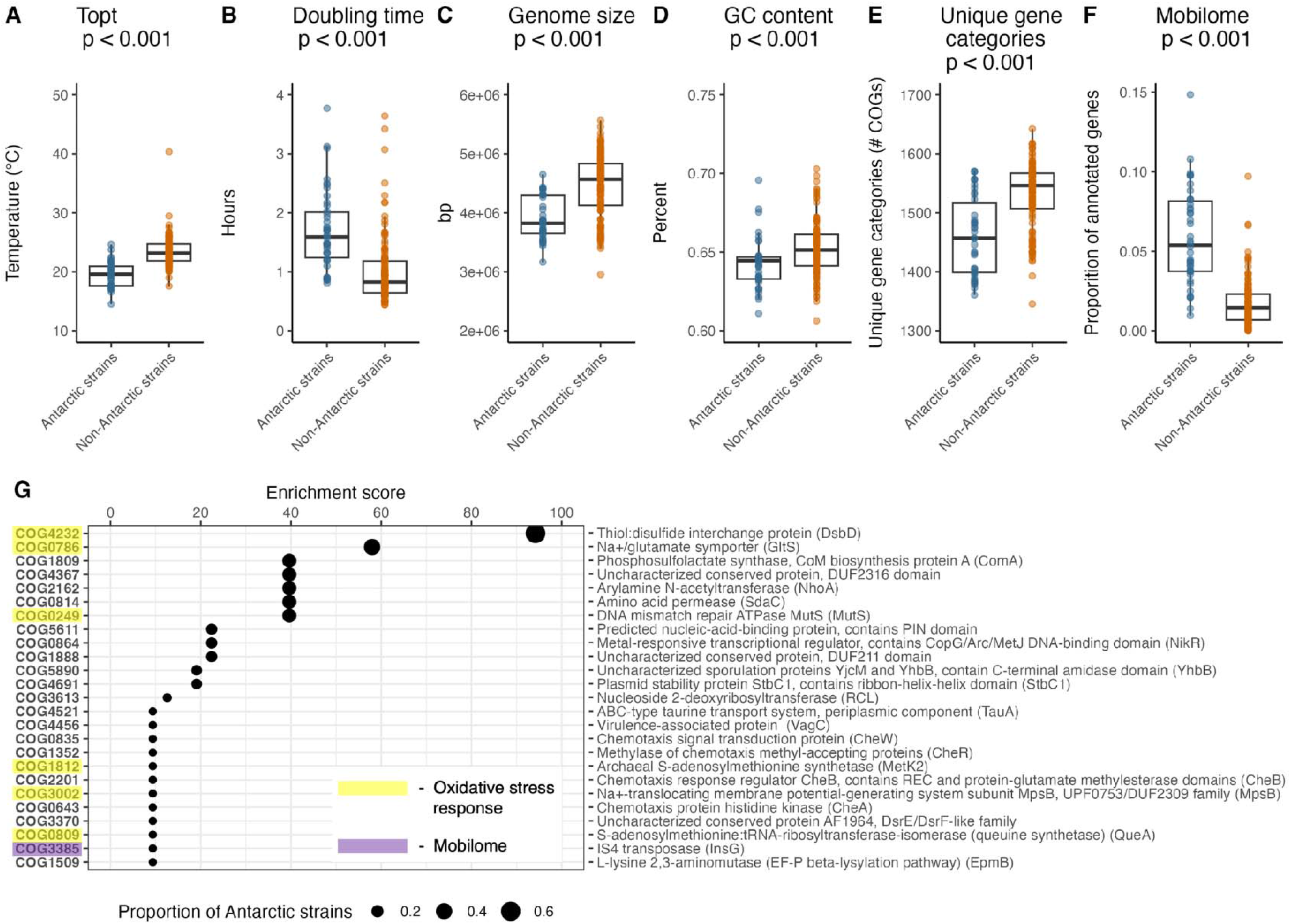
Comparison of genomic traits between Antarctic *Arthrobacter* strains (n = 42) and non-Antarctic *Arthrobacter* strains (n = 129). **A)** Estimated temperature optima (Topt) of the representative strains, as inferred using GenomeSpot (130). **B)** Estimated minimum doubling times of the representative strains, as inferred using gRodon2 (54). **C)** Differences in genome sizes across Antarctic and non-Antarctic strains. **D**) Comparison of GC content between Antarctic and non-Antarctic *Arthrobacter* strains. **E)** Differences in the number of unique gene categories (COGs) identified in the Antarctic and non-Antarctic strains. **F)** Comparison of the proportion of annotated genes associated with the mobilome (COG X) between the Antarctic and non-Antarctic strains. **G)** Genes enriched in Antarctic *Arthrobacter* strains. COGs shown here were identified as being significantly enriched (p < 0.05) in Antarctic strains, based on metrics described in ref (129). The enrichment score - Rao test statistic for equality of proportions, for each COG is represented on the X axis, while the size of the dot represents the proportion of Antarctic strains (n = 46) that contain that COG. For each COG, a more detailed annotation of the gene represented is displayed to the right of the chart. COGs are highlighted based on whether they have previously been reported to be involved in oxidative stress response or the mobilome

One characteristic feature of Antarctic soils, besides being very cold and dry, is that they typically have very low levels of organic carbon (30, 53). As a result, we would expect endemic Antarctic *Arthrobacter* strains to be more oligotrophic than strains from non-Antarctic soils (34). Our results strongly support this hypothesis. We found that the genome-derived estimates of minimum doubling times (a metric of oligotrophy (34, 54)) were significantly longer for Antarctic strains compared to non-Antarctic strains (Mann-Whitney U, p < 0.001, Figure 4), which is consistent with our growth assay results (see above and Figure 3). We also found that Antarctic strains had significantly smaller genomes (Mann-Whitney U, p <0.001, Figure 4) with fewer unique gene categories than non-Antarctic strains (Mann-Whitney U, p < 0.001, Figure 4). Smaller genome size is often presumed to be a trait associated with oligotrophic bacteria living in challenging environments, reflecting reduced metabolic costs (55–57). We also found that Antarctic strains had significantly fewer genes associated with energetically expensive activities, including cellular motility (COG N), intracellular transport and metabolisms (COGs E,I,G), and cell wall biogenesis (COG M) (Figure 4); general genomic features considered typical of bacteria with oligotrophic life history strategies (58–60).

The genomic analyses revealed certain genes and gene categories that were significantly enriched in Antarctic *Arthrobacter* strains (Figure 4, S4). Some of these have previously been linked to microbial survival across the southern continent, including genes associated with the generation of energy from diverse sources (e.g. sulfide oxidation genes and bacteriorhodopsin, Figure S5) that were enriched in Antarctic genomes (61). Unexpectedly, we did not find genes associated with trace gas metabolism to be enriched in Antarctic strains (35, 61, 62). One potential explanation for this is that these genes may also confer survival in other cold, dry environments (e.g, Atacama Desert, Arctic soils, alpine environments) which were represented in our non-Antarctic category (63). Instead, the most differentially abundant gene category in the Antarctic strains was COG category X – Mobilome (64, 65) which was nearly 300% more abundant in Antarctic strains on average (Figure 4). Enrichment of genes associated with mobile genetic elements can increase rates of horizontal gene transfer and generate the genetic variation required for organisms to adapt to environmental change (66–69). The higher abundance of these genes which facilitate the movement of DNA segments (70), including transposases (i.e., COG3385; Figure 4), may be an indicator of the evolutionary processes that have led to the divergence of Antarctic *Arthrobacter* strains from the other strains in non-Antarctic soils (71). In Antarctic soils, where population heterogeneity is expected to be relatively low and generation time is expected to be slow, speciation and evolution may be driven more by horizontal gene transfer than by spontaneous mutation.

Finally, the most enriched COG in Antarctic *Arthrobacter* strains was COG4232 (Figure 4). This gene encodes for the transmembrane protein reductase thiol:disulfide interchange protein DsbD, which facilitates disulfide bond formation (72, 73). Its enrichment could serve to increase the rate of protein folding and protein stability in cold environments (74). However, DsbD also plays a role in combating oxidative stress. Reactive oxygen species (ROS) production is high is Antarctica due high UV exposure and extreme cold, stressors which may trigger the production of these compounds as a response to damage (75, 76). DsbD helps maintain redox balance within cells by transferring reducing agents from the cytoplasm to the periplasm (77). Maintaining intracellular redox balance in this way can prevent oxidative damage, which otherwise would inhibit growth at low temperatures (78–81). While DsbD specifically has not previously been linked to survival in Antarctic conditions, other genes associated with maintaining cellular redox homeostasis (including COG0786, COG0249, COG18112, COG3002, COG0809) were also found to be enriched in Antarctic strains (Figure 4), supporting previous work suggesting that these gene categories may be particularly critical for bacterial survival in cold and dry environments (82–87).

## Conclusions

### Antarctic *Arthrobacter* are unique

First, we found limited overlap between Antarctic and non-Antarctic *Arthrobacter* strains despite both Antarctic and non-Antarctic soils harboring large amounts of phylogenetic diversity within the genus (Figs. 1, 2), suggesting that a majority of strains appear to be endemic to Antarctica. Second, we found that Antarctic *Arthrobacter* strains have unique environmental tolerances and growth dynamics compared to those found in non-Antarctic soils, specifically lower optimal growth temperatures and lower maximum growth rates (Figure 3), which suggest local adaptation to the Antarctic environment. These phenotypic results from isolates were supported by our genomic comparisons (Figs. 4) which showed that Antarctic strains are more likely to have genomic features associated with oligotrophy compared to their non-Antarctic counterparts, longer estimated minimum doubling times, smaller genomes, and an enrichment in genes associated with horizontal gene transfer and protection against oxidative damage. Together our results suggest that both geographic isolation and environmental uniqueness, factors that are known to contribute to endemism in other taxa (9, 19, 88), likely contribute to the apparent endemism of Antarctic *Arthrobacter*.

We are neither the first group to find evidence of microbial endemism in Antarctica (19, 88, 89), nor the first to propose that *Arthrobacter* in Antarctic soils may be distinct from those found in other global soils (36). Our analysis builds on this previous work, confirming that signals of endemism and local adaptation can be measured in phylogeny, environmental tolerances, and genomic traits. We were able to identify these distinctions in a globally ubiquitous bacterial genus (37, 38), highlighting the utility of the strain-level approach. Consequently, we suspect that there are other Antarctic microorganisms with comparable levels of endemicity and uniqueness, as has been previously proposed (19). Evidence for microbial endemism in Antarctica holds particular significance for the conservation of this continent and warrants careful consideration. Given that ecosystems in Antarctica are shifting rapidly due to the impacts of climate change (90, 91) and other anthropogenic pressures, including the introduction of non-indigenous microbes into fragile Antarctic environments (92), we need to investigate and identify other Antarctic microorganisms that are unique to the continent before it is too late.

## Materials and Methods

Strain-level biogeographic analyses from shotgun metagenomes: First, we compiled shotgun metagenomic datasets from a wide range of Antarctic and non-Antarctic soils to characterize the diversity and distributions of members of the *Arthrobacter* genus to determine if the strains detected in Antarctic soils are shared with those found elsewhere. For the Antarctic soil metagenomes, we combined 27 soil metagenomes from ref (29) (NCBI SRA accession: PRJNA699250) with metagenomic data from an additional 99 Antarctic soil samples that were collected as described in ref (30). These samples were chosen from the larger set of samples described in ref (30) as they had sufficiently high DNA concentrations (>5 ng/ul). Metagenomic sequencing largely followed methods described in (29, 93).

Briefly, DNA aliquots from each sample were used to generate a metagenomic library with the Nextera DNA Flex library preparation kit (Illumina, San Diego, CA, USA). This library was sequenced on two lanes of an Illumina NovaSeq 6000 using a high output 300-cycle kit with paired-end chemistry at Texas A&M’s Genomic and Bioinformatics Service Center. The raw metagenomic data has been deposited in the NCBI Sequence Read Archive, project accession number PRJNA1214840. Together, the 126 Antarctic soil metagenomes used for this study span a wide range of soil types and locations across the continent (Figure 1).

We next compiled a total of 472 metagenomes from soils found outside Antarctica for comparison (Figure 1). These non-Antarctic soil metagenomes included: 331 surface soil metagenomes from the Biomes of Australian Soil Environments (BASE) Project (94, 95) (NCBI accession PRJNA597010), 15 soil metagenomes generated by Brewer et al. (96) from surface soils collected as part of the Critical Zone Observatory Network (MG-RAST project ID mgp80869), 32 soil metagenomes generated by the National Ecological Observatory Network, release 2023 (available on the Joint Genome Institute (JGI) data portal (https://data.jgi.doe.gov/) under JGI_ID 509938) (97), 41 soil metagenomes collected across China, and 53 soil metagenomes from cold-dry environments across the Arctic (Greenland, Svalbard, Alaska, Canada) (98–100), alpine regions in Europe and across the Tibetan plateau (100, 101), and the Atacama desert (JGI project 3300061561). We selected these datasets as the metagenomes were publicly available and were all exclusively derived from surface soils that span a wide range of unmanaged ecosystem types, with the sample collection, processing, and metagenomic data generation conducted using methods similar to those used for the 126 Antarctic soil metagenomes described above. Mean annual temperature (MAT) and mean annual precipitation (MAP) for each soil were determined following the methods described in ref (30). Briefly, MAT (×C) and MAP (mm/year) were extracted from the CHELSA database v.2.1 (102) using CHELSA variables BIO1 and BIO12 (1981 – 2010) respectively. MAT and annual precipitation could not be calculated for 5 metagenomes where precise GPS locations were lacking. Information about each metagenome, including NCBI BioProject and BioSample IDs can be found in Dataset S1.

We utilized a strain-level approach (103) to analyze *Arthrobacter* distributions across the 598 unique surface soil metagenomes (126 from Antarctica and 472 from outside Antarctica). We first quality filtered the metagenomes with Trimmomatic v.0.39 (104). We then used the “profile” function of sylph v.0.8.1 (105) to search for a reference set of 568 *Arthrobacter* genomes across each these 598 soil metagenomes. We define strains here as members of the genus that have distinct genomes (103). The reference set of *Arthrobacter* genomes included the assembled genomes of our 53 isolates (described below) and 515 high quality (completeness >95, contamination <5% (106)) *Arthrobacter* genomes and metagenome-assembled genomes (MAGs) from GTDB v.220 (107). Sylph presence and absence results can be found in Dataset S2.

Isolation and genome sequencing of soil *Arthrobacter* strains: To complement the metagenomic analyses, we also used a cultivation-dependent approach, isolating a total of 53 unique *Arthrobacter* strains from Antarctic and non-Antarctic soils. A total of 28 unique *Arthrobacter* isolates from Antarctic soils were selected from the culture collection described in ref (30). Non-Antarctic *Arthrobacter* isolates were chosen from several studies: 8 isolates were from soils collected from the former volcanic island of Hunga Tonga Hunga Ha’apai (93), 2 isolates were from soils from the Mahalangur Himal sub-range of the Himalayas in Nepal (49), 1 isolate was from a tropical forest soil in Panama(108), 1 isolate was from a soil collected in Australia (17), and 8 isolates from soils collected at Harvard Forest, MA, USA (109). Finally, 5 *Arthrobacter* type strains were purchased that were originally isolated from soils in Austria (DSMZ 22274), the United Kingdom (DSMZ 14008), South Korea (DSMZ 16760), China (DSMZ 24664), and Japan (DSMZ 25586). The 53 isolates used in our study were re-grown from glycerol stock on Reasoners 2 Agar (MilliporeSigma, Haward, CA, USA) and were re-streaked at least three times to ensure purity. More details on the *Arthrobacter* isolates used in this study can be found in Dataset S3.

To confirm isolate identity, we performed full length 16S rRNA gene sequencing and whole genome sequencing. DNA was extracted from the 53 cultures with the Qiagen DNeasy Ultraclean Microbial kit (Qiagen, Redwood City, CA, USA) following manufacturers recommendations. The DNA aliquots from each of the bacterial isolates were then PCR amplified using primer pairs that target the 16S rRNA gene (8F: 5’-AGAGTTTGATCCTGGCTCAG-3’ and 1391R: 5′-GACGGGCGGTGWGTRCA -3′) (110, 111). PCRs were performed in 25µL-reaction volumes using PlatinumTM II Hot-Start PCR Master Mix (2X) (Invitrogen, Carlsbad, CA, USA) on a SimpliAmp Thermal Cycler (Thermo Fisher Scientific, Waltham, MA, USA). Cycling parameters for the primer set consisted of an initial denaturation step at 94 °C for 2 minutes, followed by 35 cycles of denaturation at 94 °C (15 s), annealing at 60°C (15 s), extension at 68 °C (60 s), and a final extension step at 72 °C for 10 minutes. All amplified PCR products were sequenced with the forward 8F primer by Azenta Life Sciences’ Sanger sequencing service (Azenta Life Sciences, Burlington, MA, USA). Sequences were processed following a modified “Swabs to Genomes” workflow described in refs (49, 112). All sequences >1000bp in length were cleaned with SeqTrace v.0.9.1 (113) after which taxonomy was assigned using the RDP classifier v.2.10.2(114). 16S rRNA gene sequences for all 53 isolates are reported in Dataset S4.

Aliquots of the extracted DNA from each isolate were also prepared for whole genome sequencing with the Oxford Nanopore Native Barcoding Kit 96 V14 (Oxford Nanopore, Oxford, UK). Sequencing was performed on a MinION Mk1B (Oxford Nanopore, Oxford, UK) with a R10.4.1 flow cell and kit 14 chemistry (ONT). Base calling and demultiplexing were performed with Dorado (https://github.com/nanoporetech/dorado), following the “super accurate” model. The raw sequencing data was filtered with Filtlong v.0.2.1 (https://github.com/rrwick/Filtlong) following default parameters to remove the 5% lowest quality reads. Filtered reads were then assembled with Flye v.2.9.5 (115) with parameters for high quality ONT reads. Additional binning of assemblies was performed with MaxBin2 v.2.2.7 (116). The generated assemblies were polished with Medaka v.2.0.1 (https://github.com/nanoporetech/medaka) and Polypolish v.0.6.0(83, 117). Assembly quality was assessed with CheckM2 (118), open reading frames were predicted with Prodigal v.2.6.3 (119), and taxonomy was assigned with the GTDBtk v.2.1.0 classify workflow (120) based on the Genome Taxonomy Database v.220 (107). Average nucleotide identify (ANI) between all genomes was calculated with FastANI (121). We note that all 53 isolates were confirmed to be members of the *Arthrobacter* genus with each isolate representing a unique strain: mean average nucleotide identity of 83.9%, range 77.8 – 99.9%.

### Strain-level genomic comparisons

We analyzed our *Arthrobacter* genomes to compare functional attributes between Antarctic and non-Antarctic strains. These analyses were conducted with genomic data from *Arthrobacter* strains found in Antarctic soils: 32 strains detected by sylph during the metagenomic profiling of the Antarctic soils described above and 10 additional strains from the isolates obtained as described above. For the strains identified from non-Antarctic soils, we included genomic data from the 107 strains detected by sylph in non-Antarctic soils from our metagenomic analyses and 21 additional strains isolated from non-Antarctic soils as described above. In total, the genomic analyses were based on 42 unique strains of *Arthrobacter* from Antarctic soils and 129 unique *Arthrobacter* strains from non-Antarctic soils. The genomes from all 171 strains included in the genomic analyses were all considered to be high quality (>95% complete with <5% contamination (106) (see Dataset S4 for more details).

Phylogenetic analyses of the 171 strains were conducted using whole-genome data from each strain. Multiple sequence alignments (MSA) of protein sequences from *Arthrobacter* genomes were created using the GTDB-Tk “align” function (--skip_gtdb_refs --custom_msa_filters --cols_per_gene 3000 -- min_consensus 25 --max_consensus 95 --min_perc_taxa 100) (120). We used more amino acids per gene than the default settings but only included proteins that were present in all 171 strains, resulting in a total of 2000 amino acids used in the alignment. FastTree v.2.2 (122) was then used to infer approximately-maximum-likelihood phylogenetic trees from the MSA, with GTDB genome GCF_000725885.1 (*Streptymyces albus*) included as the outgroup. Trees were visualized and annotated using iTOL v.7.2 (123).

To compare the genomic attributes of the strains from these two categories (Antarctic vs. non-Antarctic), we followed a modified version of the Anvi’o pan-genomic workflow v.7.1 (124). Briefly, open reading frames were identified with Prodigal (119) amino acid sequence similarities across our 171 *Arthrobacter* genomes were calculated with Diamond (125), weak matches between sequences were excluded with ITEP (126), clusters of similar amino acid sequences were identified with the MCL algorithm (127), and Euclidean distance and Ward linkage clustering were used to organize gene clusters and identify similarity across genomes. Genes were annotated using the Anvi’o “anvi-display-functions” workflow and the Cluster of Orthologous Genes (COG) database v.2024 (65) and the Protein Families database (Pfam) (128). Functional enrichment analysis comparing Antarctic and non-Antarctic strains was performed with the “anvi-compute-functional-enrichment” program and exported with the “anvi-script-gen-function-matrix-across-genomes” function (124), which calculates enrichment scores based on the methods described in ref (129). Additional genomic features were collated from the results of GTDB-TK and CheckM2 (summarized previously). Pairwise ANI comparisons were performed with FastANI v.1.34 with the default cutoff for inclusion (∼77%) (121). We used the tool gRodon2 to estimate maximum potential growth rates from codon usage biases in highly expressed genes (54) following the workflow described in ref (29). We used GenomeSpot v.1.0.1 (130) to predict optimal growth temperature, minimum growth temperature, and maximum growth temperature from the genomes. Finally, additional genomic features were identified using the methods described previously (29, 93, 131) Briefly, we used the blastp function of Diamond v. 2.1.16 (125) to search for the 52 metabolic marker genes in a custom protein database (61, 131) (https://github.com/GreeningLab/GreeningLab-database) consisting of marker genes involved in carbon fixation, nitrogen cycling, phototrophy, respiration, sulfur cycling, trace gas metabolism, and alternate electron acceptors and donors.

Statistical comparisons were performed in R v.4.3.2 (132) with the package ‘rstatix’ (https://github.com/kassambara/rstatix). Homogeneity of variance assumptions were tested with Levene’s tests and normality of distribution was calculated with Shapiro-Wilk tests. To identify differences in the genomic characteristics, predicted maximum growth rates, and abundance of COG categories between the 42 Antarctic strains and the 129 non-Antarctic strains, we used Mann-Whitney nonparametric tests corrected for multiple comparisons with Bonferroni tests. Comparisons made across more than three groups were performed with Kruskal-Wallis tests with post-hoc Nemenyi tests to calculate pairwise multiple comparisons between groups. Additional dissimilarity measurements were performed with the R packages ‘mctoolsr’ (https://github.com/leffj/mctoolsr/), ‘vegan’ (https://github.com/vegandevs/vegan), and ‘phyloseq’ (133). ‘Picante’ was used to calculate Faith’s phylogenetic diversity (134). Plotting was performed with the packages ‘ggplot2’ (135) and ‘cowplot’ (136). Maps were created in QGIS v.3.44 (137) with the “QuickMapServices” plugin for the world map. We used Quantarctica v.3.2 (138) for the Antarctic map using the “Basemap” and “SimpleBasemap” data layer which used the SCAR Antarctic Digital Database (ADD) v.7.0. Downloaded 2016-2017 from: http://www.add.scar.org/. Coastline polygons north of 60°S were taken from 1:10000000 datasets available on Natural Earth (http://www.naturalearthdata.com/) v.3.0.0.

To test whether the number of genomes that were detected in both Antarctic and non-Antarctic metagenomes was lower than expected by chance, we generated a null distribution by randomly assigning samples as Antarctic or non-Antarctic 1000 times (while maintaining n = 50 Antarctic samples and n = 105 non-Antarctic samples) and each time calculated the number of genomes detected in both sample types. We then ran a hypergeometric Test (equivalent to a one-tailed Fisher’s Exact Test) based on a total population size of (N=147) to test whether the observed overlap (k=4) between Antarctic strains (Group A, K=36) and non-Antarctic strains (Group B, n=111) was statistically significant or attributable to random chance. Analyses were performed in R (132) using ‘Vegan’ and ‘phyper’ to calculate hypergeometric distribution significance (139). Statistical significance was defined at p = 0.05.

### Isolate Growth Assays

We used the 53 unique *Arthrobacter* isolates described above (28 from Antarctic soils, 25 from non-Antarctic soils) to quantify their growth responses across gradients in water availability and temperature. For the water availability and temperature assays we used media infused with varying concentrations of polyethylene glycol 6000 (PEG, ThermoFisher scientific, Pittsburgh, Pennsylvania, USA) following methods modified from refs (81, 140–143) to mimic a gradient in matric potential where media with higher PEG concentrations have lower (more negative) water potentials and higher matric stress. Briefly, R2A agar with pH adjusted to 7.2 using 1M HCl was prepared with 2% agar. The media was then autoclaved on a liquid cycle with a 45-minute sterilization. One mL of the R2A solution was then poured into each well of a 24 well Corning Costar Plate (Corning Inc., Corning, NY, USA) and left to solidify. Five different concentrations of PEG (0% - 250% PEG) were then prepared by dissolving PEG in a Tris(hydroxymethyl)aminomethane (Tris) buffer solution (MP Biomedicals, Burlingame, CA,), with pH adjusted to 7.2 using 1M HCl. The 0% PEG solution was prepared following the same methods, but without any PEG added. PEG solutions were then filter sterilized with 0.2 µM vacuum filtration unit (ThermoFisher Scientific, Waltham, MA, USA). 1.5 mL of each PEG solution were pipetted on top of the R2A media in the wells of the plates so that each column (4 wells) had the same concentration of PEG, and concentration increased in each row from left (0% PEG) to right (250% PEG). PEG was left to infuse in the R2A media for 24 hours after which excess PEG was poured off.

The 53 *Arthrobacter* isolates (28 Antarctic, 25 non-Antarctic) were plated on 5 replicate 24-well plates prepared with PEG solution. Each isolate was spotted into the center of each well, creating 4 replicates of each culture at each PEG concentration per plate. Plates were then incubated at five different temperatures (4ºC, 11ºC, 18ºC, 25ºC, 32ºC) for seven days. After seven days, plates were photographed and then frozen. Colony growth from all wells on all plates (120 measurements per culture) was measured from these photographs with ImageJ (144).

Temperature optima were predicted for each culture with generalized additive models (GAMs). Models were built using the day seven growth measurements for the 0% PEG replicates across the 5 temperatures (4 replicates/temperature, 20 total). Reported temperature optima are the predicted temperatures at which grow of each *Arthrobacter sp*. was maximized. We used the gam.check function of the r package ‘mgcv’ (145) to calculate diagnostics for each model. Models included in this analysis had a pval < 0.05 with a k-index > 0.5. Parameters, including k’, effective degrees of freedom, and k-index can be found in Dataset S5.

Desiccation tolerances were also modeled using GAMs as above. Models used the measurements from the 24 replicates (4 per PEG concentration) on the plate grown at the temperature closest to the predicted temperature optima (See Dataset S5). We used two metrics of desiccation tolerance: the predicted PEG concentration at which growth was reduced by 50% from maximum growth, and the fold change in growth of each culture from the average growth at 0% PEG (highest water availability). Reported water potential (MPa) was determined from the concentration of PEG in each of our solutions (141, 146).

## Acknowledgments

We thank the members of the Fierer and Quandt Labs at CU Boulder for help and suggestions with the laboratory analyses and nanopore sequencing. We also thank Profs. Craig Cary, Diana Wall, Leopoldo Sancho, Charles K. Lee, John E. Barrett for collecting and providing the soil samples. We also acknowledge the Norwegian Polar Institute and Quantarctica, whose data we used to create the Antarctic map in Figure 1, and Texas A&M’s Genomic and Bioinformatics Service Center, Plasmidsaurus, and Azenta Life Sciences for their sequencing services. Funding was provided by US National Science Foundation Office of Polar Programs grant NSF 21-567 (NF, CAQ, BA), Australian Research Council grant FT240100502 (CG), Australian Research Council grant DE250101210 (PML), US. National Science Foundation grant EAR 1950681 (salary for NM)

## Data, code, and materials availability

All data used in this study are available in the main text or the supplementary material. The metagenomic sequencing data we generated from our 99 Antarctic soils and the whole genome sequencing data from our 53 Arthrobacter isolates are available in the National Center for Biotechnology Information Sequence Read Archive: BioProject ID PRJNA1214840. All other metagenomes used in the analysis are publicly available from previous studies, with repository and accession information presented in Dataset S1. Additional information about the soil samples can be found in the Environmental Data Initiative (EDI) Repository DOI:10.6073/pasta/b4858653c587864f0111aba4c3014d61 with more detailed information on the isolates available on Figshare: https://doi.org/10.6084/m9.figshare.28266821.v1.

## Notes

### Competing Interest Statement

The authors have declared no competing interest.

