## Supplemental Information for "Evidence for endemism and local adaptation in Antarctic soil bacteria"

**This PDF file includes:**

Figures S1 to S5

Legends for Datasets S1 to S5

SI References

**Other supporting materials for this manuscript include the following:**

Datasets S1 to S5

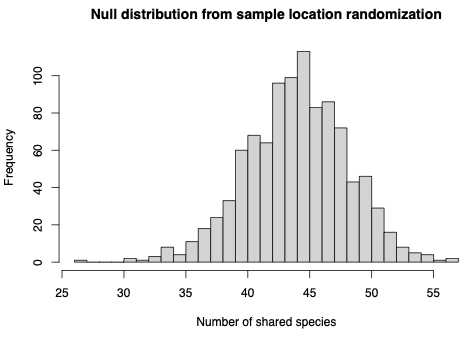

**Figure S1.** Display of the null distribution created by randomly assigning samples as Antarctic or non-Antarctic 1000 times (while maintaining n = 50 Antarctic samples and n = 105 non-Antarctic samples). The histogram displays the frequency that each number of genomes was detected in both sample types”

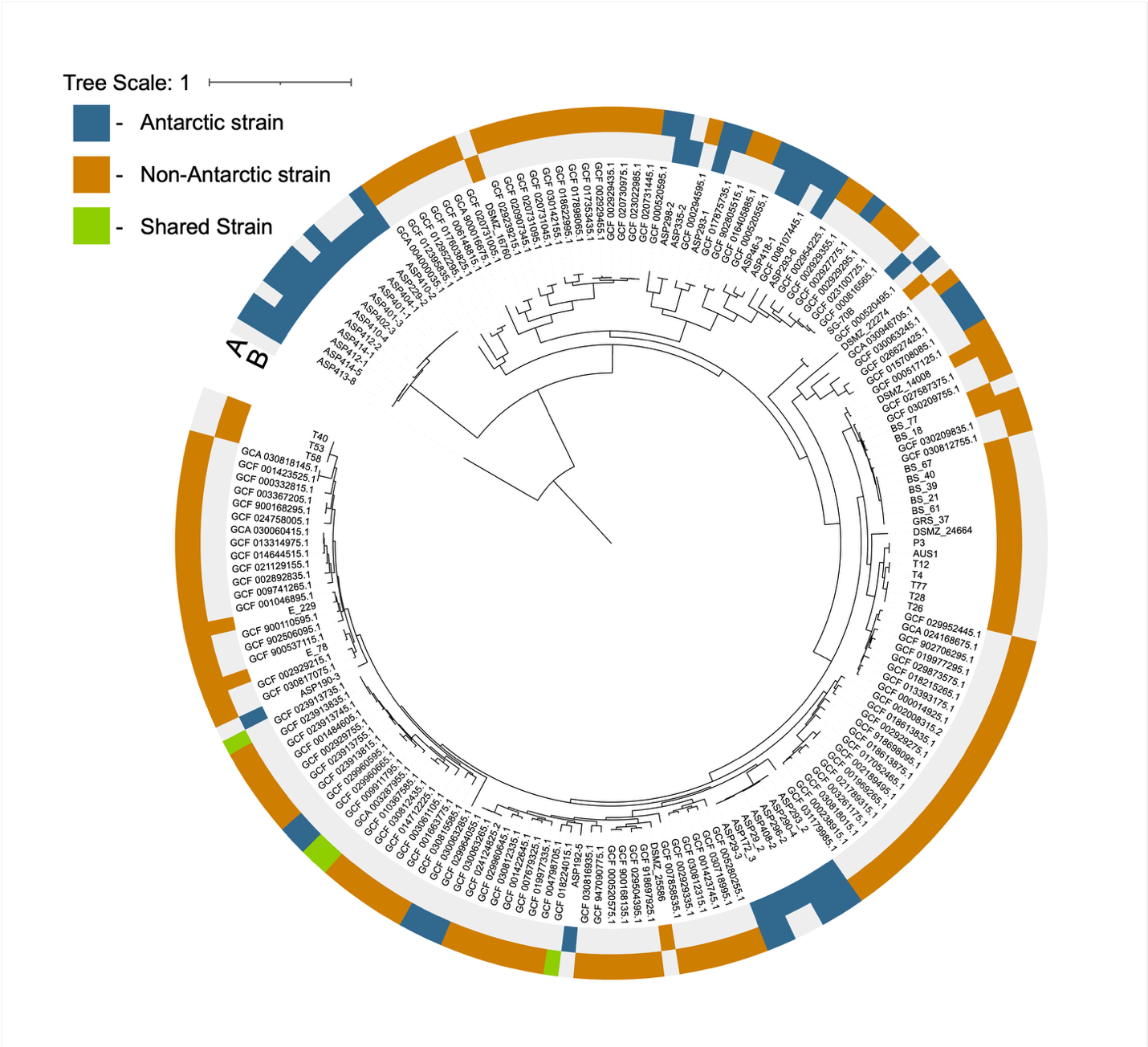

Figure S2. Phylogenetic tree of the 42 Antarctic *Arthrobacter* strains, the 129 non-Antarctic *Arthrobacter* strains, and four shared strains. The strains represented in this tree include all isolates from Antarctic and non-Antarctic soils (See methods for more details) and all strains identified through metagenome profiling (see Fig. 1). Strains are labeled either with the genome ID or isolate ID. The outer ring (“A”) indicates the strains that were identified through metagenome profiling, while the inner ring (“B”) indicates whether the strains were directly isolated from Antarctic or non-Antarctic soils. Strains associated with Antarctica are highlighted in blue, non-Antarctic strains are highlighted in orange, shared strains are highlighted in green.

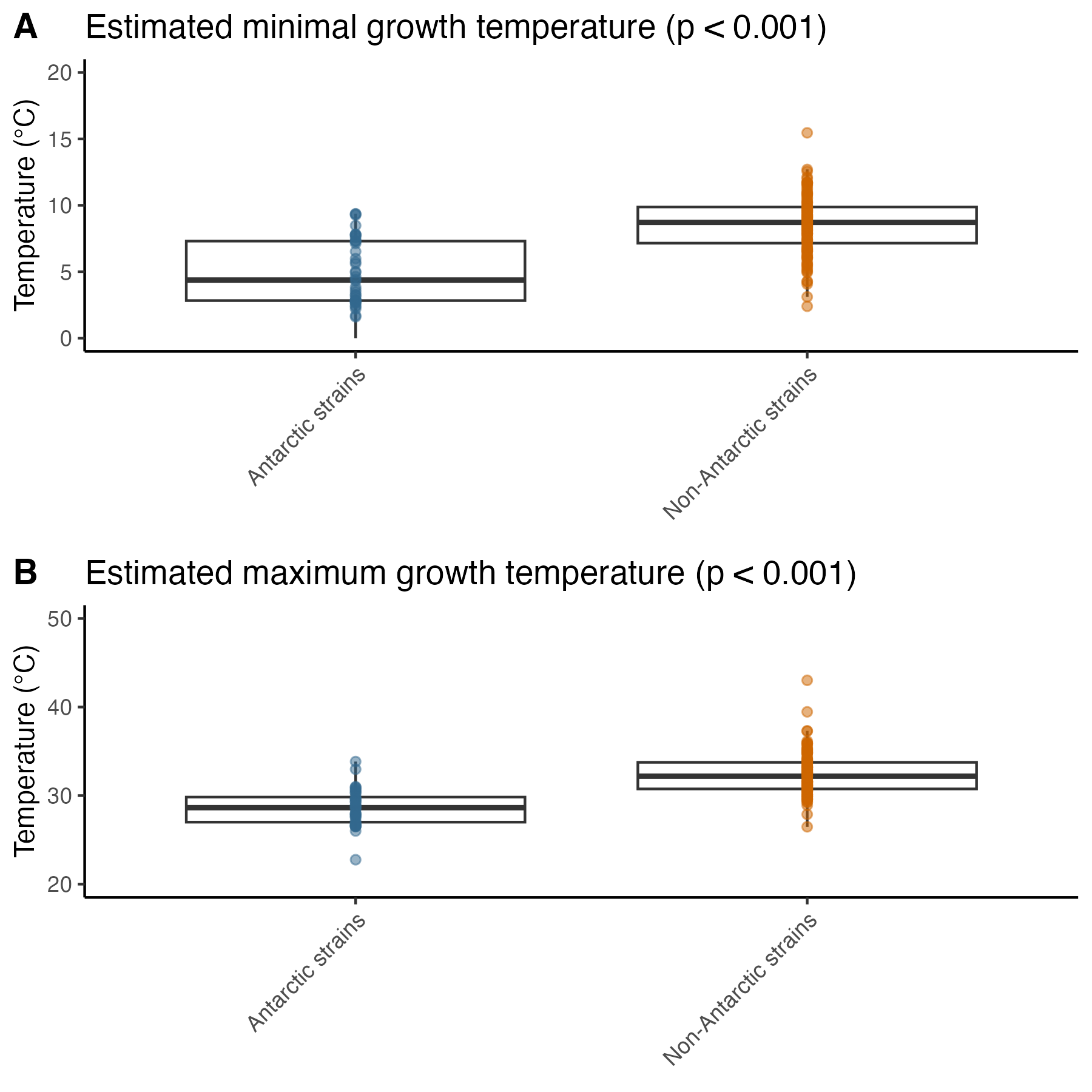

Figure S3. Comparison of estimated growth rates of the Antarctic (n=42) and non-Antarctic strains (n=129) calculated with GenomeSpot (1)*.*  A) Estimated minimum growth temperatures of Antarctic and non-Antarctic strains. B) Estimated maximum growth temperatures of Antarctic and non-Antarctic strains. Significance (Mann-Whitney U, pval), is highlighted above each plot.

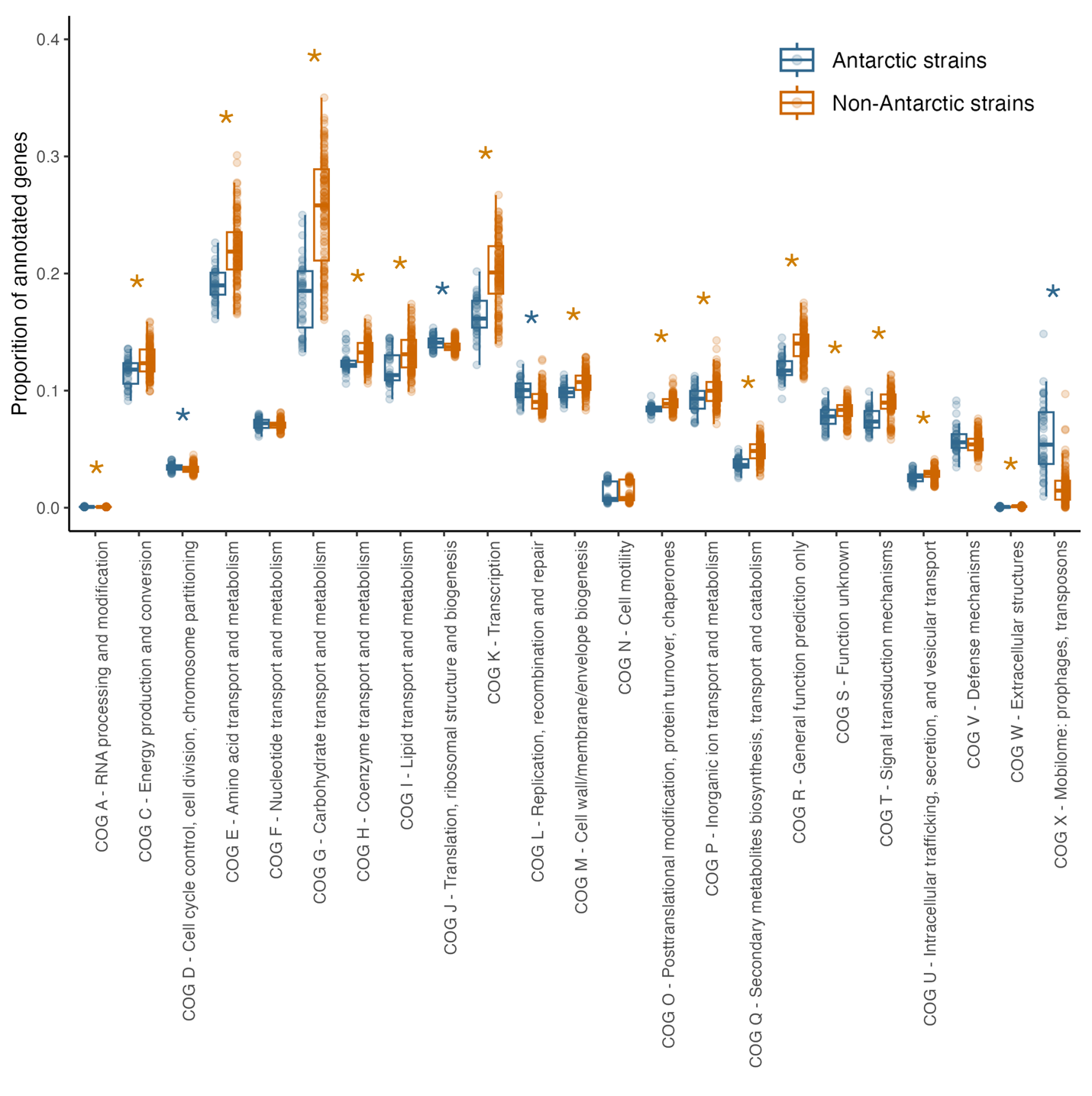

Figure S4. General genomic attributes of Antarctic *Arthrobacter* strains (n = 42) and non-Antarctic *Arthrobacter* strains (n = 129). Comparison of the proportion of annotated genes associated with each COG category between the Antarctic and non-Antarctic strains. A significant difference in a category (Mann-Whitney U, p < 0.05) is highlighted by an asterisk with the color indicating the group (Antarctic or non-Antarctic) whose genomes contain a higher proportion of genes associated with that category.

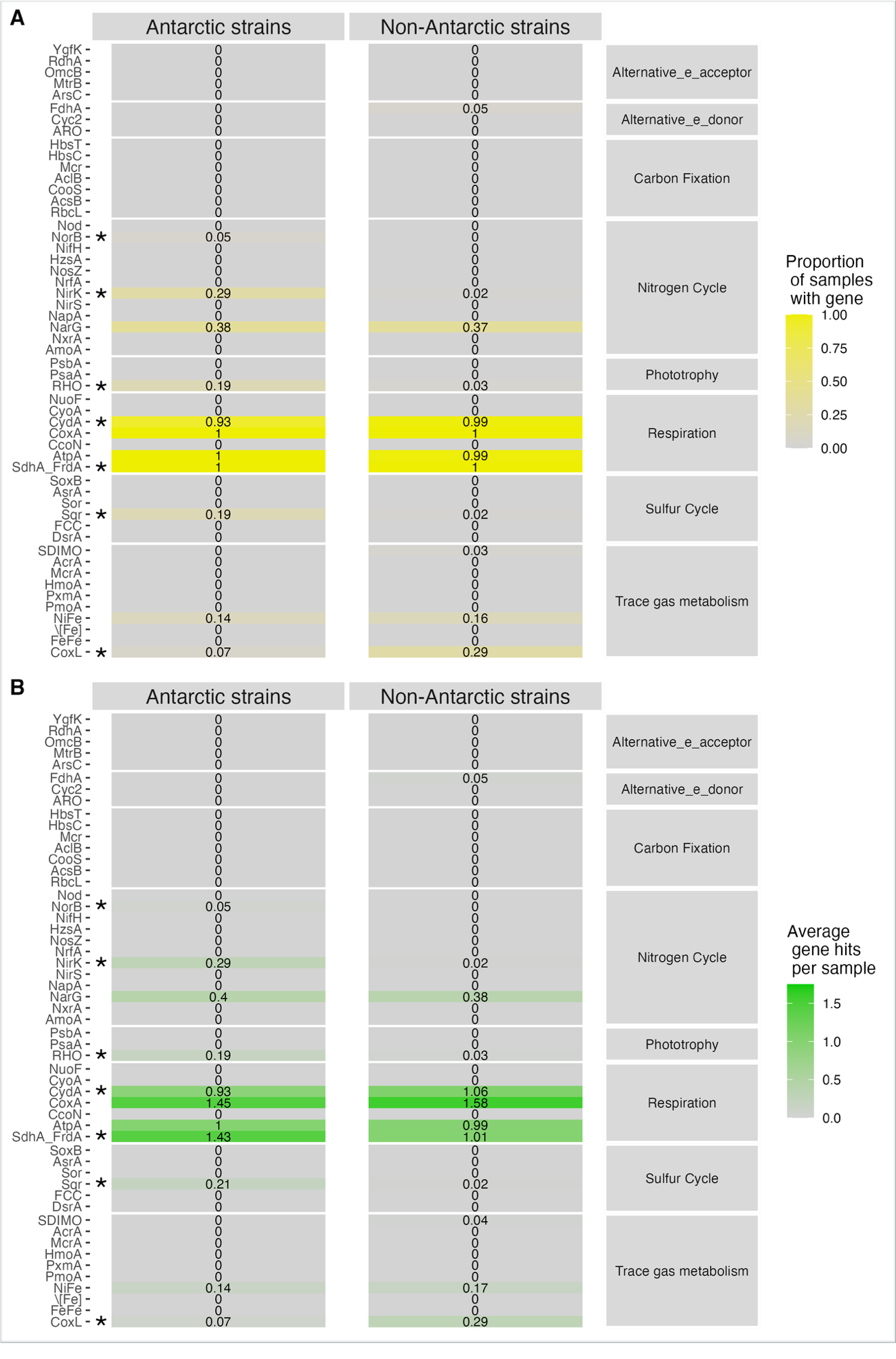

Figure S5. Abundance of functional genes in the genomes of Antarctic *Arthrobacter* strains (n = 42) and non-Antarctic *Arthrobacter* strains (n = 129). A) Proportion of samples within each category (Antarctic strain vs. non-Antarctic strain) containing at least one hit to the 52 genes surveyed in our gene search (see methods for more details). B) Comparison of the average number of gene hits per sample for each category. A significant difference in the two sample groups category (Mann-Whitney U, p < 0.05) is highlighted by an asterisk next to the gene name in each plot. The 53 genes are grouped into eight larger functional categories, which are displayed to the right.

Dataset S1. (separate file Arthrobacter_Dataset_S1.xlsx)

This file contains the information about the 126 Antarctic soil metagenomes and 472 non-Antarctic metagenomes used in our Sylph metagenomic profiling. Details include NCBI or JGI accession, sample ID, and predicted mean annual temperature (MAT) and mean annual precipitation (MAP).

Dataset S2. (separate file Arthrobacter_Dataset_S2.xlsx)

This file contains the presence and absence table of *Arthrobacter* genomes identified by Sylph during our metagenomic profiling. If a genome was identified in a sample, it is indicated with a “1” otherwise “0” indicates that genome was not identified in that sample.

Dataset S3. (separate file Arthrobacter_Dataset_S3.xlsx)

This file contains the information about the 53 *Arthrobacter* isolates used in the cultivation independent analyses, including: details about the quality and completeness of each genome and the 16S rRNA gene sequence.

Dataset S4. (separate file Arthrobacter_Dataset_S4.xlsx)

This file contains the results of the genomic analysis comparing the 42 Antarctic strains and the 129 non-Antarctic strains. Results contained include CAzyme abundance, gRodon maximum potential growth rate, predicted temperature optima, as well as all COG categories and COG abundances.

Dataset S5. (separate file Arthrobacter_Dataset_S5.xlsx)

This table contains the raw data from the temperature and desiccation assays as well as the results and metrics of the generalized additive models used to predict temperature optima and desiccation tolerance.
